# Genome-wide association analyses identify variants in *IRF4* associated with acute myeloid leukemia and myelodysplastic syndrome susceptibility

**DOI:** 10.1101/773952

**Authors:** Junke Wang, Alyssa I. Clay-Gilmour, Ezgi Karaesmen, Abbas Rizvi, Qianqian Zhu, Li Yan, Leah Preus, Song Liu, Yiwen Wang, Elizabeth Griffiths, Daniel O. Stram, Loreall Pooler, Xin Sheng, Christopher Haiman, David Van Den Berg, Amy Webb, Guy Brock, Stephen Spellman, Marcelo Pasquini, Philip McCarthy, James Allan, Friedrich Stölzel, Kenan Onel, Theresa Hahn, Lara E. Sucheston-Campbell

## Abstract

The role of common genetic variation in susceptibility to acute myeloid leukemia (AML), and myelodysplastic syndrome (MDS), a group of rare clonal hematologic disorders characterized by dysplastic hematopoiesis and high mortality, remains unclear. We performed AML and MDS genome-wide association studies (GWAS) in the DISCOVeRY-BMT cohorts (2309 cases and 2814 controls). Association analysis based on subsets (ASSET) was used to conduct a summary statistics SNP-based analysis of MDS and AML subtypes. For each AML and MDS case and control we used PrediXcan to estimate the component of gene expression determined by their genetic profile and correlate this imputed gene expression level with risk of developing disease in a transcriptome-wide association study (TWAS). ASSET identified an increased risk for *de novo* AML and MDS (OR=1.38, 95% CI, 1.26-1.51, P_meta_=2.8×10^-12^) in patients carrying the T allele at rs12203592 in *Interferon Regulatory Factor 4* (*IRF4*), a transcription factor which regulates myeloid and lymphoid hematopoietic differentiation. Our TWAS analyses showed increased *IRF4* gene expression is associated with increased risk of *de novo* AML and MDS (OR=3.90, 95% CI, 2.36-6.44, P_meta_ =1.0×10^-7^). The identification of *IRF4* by both GWAS and TWAS contributes valuable insight on the role of genetic variation in AML and MDS susceptibility.

## INTRODUCTION

Genome-wide association studies (GWAS) have been successful at identifying risk loci in several hematologic malignancies, including acute myeloid leukemia (AML) ^1–3^. Recently genomic studies have identified common susceptibility loci between chronic lymphocytic leukemia (CLL), Hodgkin lymphoma (HL), and multiple myeloma demonstrating shared genetic etiology between these B-cell malignancies (BCM) ^4–6^. Given the evidence of a shared genetic basis across BCM and the underlying genetic predisposition for AML and myelodysplastic syndromes (MDS) observed in family, epidemiological, and genetic association studies^1^,^7–9^, we hypothesized that germline variants may contribute to both AML and MDS development. Using the DISCOVeRY-BMT study population (2309 cases and 2814 controls), we performed AML and MDS GWAS in European Americans and used these data sets to inform our hypothesis. To address the disease heterogeneity within and across our data we used a validated meta-analytic association test based on subsets (ASSET) ^4^. ASSET tests the association of SNPs with all possible AML and MDS subtypes and identifies the strongest genetic association signal. To systematically test the association of genetically predicted gene expression with disease risk, we performed a transcriptome wide association study (TWAS)^10^,^11^. This allows a preliminary investigation into the role of non-coding risk loci, which might be regulatory in nature, that impact expression of nearby genes. The TWAS statistical approach, PrediXcan ^11^, was used to impute tissue-specific gene expression from a publicly available whole blood transcriptome panel into our AML and MDS cases and controls. The predicted gene expression levels were then tested for association with AML and MDS. The use of both a GWAS and TWAS in the DISCOVeRY-BMT study population allowed us to identify AML and MDS associations with *IRF4*, a transcription factor which regulates myeloid and lymphoid hematopoietic differentiation, and has been previously identified in GWAS of BCM.(5)

## MATERIALS AND METHODS

### Study Design & Population

Our study was a nested case-control design derived from the parent study DISCOVeRY-BMT (Determining the Influence of Susceptbility COnveying Variants Related to 1-Year Mortality after unrelated donor Blood and Marrow Transplant).^12^ The DISCOVeRY-BMT cohort was compiled from 151 centers around the world through the Center for International Blood and Marrow Transplant Research (CIBMTR). Briefly, the parent study was designed to find common and rare germline genetic variation associated with survival after an URD-BMT. DISCOVeRY-BMT consists of two cohorts of ALL, AML and MDS patients and their 10/10 human leukocyte antigen (HLA)-matched unrelated healthy donors. Cohort 1 was collected between 2000 and 2008, Cohort 2 was collected from 2009-2011.

AML and MDS patients were selected from the DISCOVeRY-BMT patient cohorts and used as cases and all the unrelated donors from both cohorts as controls. AML subtypes included *de novo* AML with normal cytogenetics, *de novo* AML with abnormal cytogenetics and therapy-related AML (t-AML). *De novo* AML patients did not have precedent MDS, chemotherapy or radiation for prior cancers. MDS subtypes included *de novo* MDS, defined as patients without precedent chemotherapy or radiation for prior cancers, and therapy-related MDS (t-MDS). Patient cytogenetic subtypes were available, however due to limited sample sizes for each cytogenetic risk group, we consider here only broad categories. Controls were unrelated, healthy donors aged 18-61 years who passed a comprehensive medical exam and were disease-free at the time of donation. All patients and donors provided written informed consent for their clinical data to be used for research purposes and were not compensated for their participation.

### Genotyping, imputation, and quality control

Genotyping and quality control in the DISCOVeRY-BMT cohort has previously been described in detail ^12–15^. Briefly, samples were assigned to plates to ensure an even distribution of patient characteristics and genotyping was performed at the University of Southern California Genomics Facility using the Illumina Omni-Express BeadChip® containing approximately 733,000 single nucleotide polymorphisms (SNPs).^16^ SNPs were removed if the missing rate was > 2.0%, minor allele frequency (MAF) < 1%, or for violation of Hardy Weinberg equilibrium proportions (P< 1.0×10^-4^).

Problematic samples were removed based on the SNP missing rate, reported-genotyped sex mismatch, abnormal heterozygosity, cryptic relatedness, and population outliers. Population stratification was assessed via principal components analysis using Eigenstrat software^17^ and a genomic inflation factor (λ) was calculated for each cohort. Following SNP quality control, 637,655 and 632, 823 SNPs from the OmniExpress BeadChip in Cohorts 1 and 2, respectively were available for imputation. SNP imputation was performed using Haplotype Reference Consortium, hg19/build 37 (http://www.haplotype-reference-consortium.org/home) via the Michigan Imputation server ^18^,^19^. Variants with imputation quality scores <0.8 and minor allele frequency (MAF) <0.005 were removed yielding almost 9 million high quality SNPs available for analysis in each cohort.

## METHODS

### Statistical Analysis

#### Genome-wide SNP associations with AML and MDS

Quality control and statistical analyses were implemented using QCTOOL-v2, R 3.5.2 (Eggshell Igloo), Plink-v1.9, and SNPTEST-v2.5.4-beta3. Logistic regression models adjusted for age, sex, and three principal components were used to perform single SNP tests of association with *de novo* MDS, t-MDS, AML by subtype (*de novo* AML with normal cytogenetics, *de novo* AML with abnormal cytogenetics and t-AML) in each cohort. European American healthy donors were used as controls. SNP meta-analyses of cohorts 1 and 2 were performed by fitting random effects models.^20^ To identify the strongest association signal with AML and MDS we conducted a summary statistic SNP-based association analysis (ASSET) implemented in R statistical software ^4^. ASSET tests each SNP for association with outcome using an exhaustive search across non-overlapping AML and MDS case groups while accounting for the multiple tests required by the subset search, as well as any shared controls between groups ^4^.

#### Heritability estimation of AML and MDS

We calculated heritability of AML and MDS combined and by independent subtypes as the proportion of phenotypic variance explained by all common genotyped SNPs, using the genome-based restricted maximum likelihood method performed with the Genome-wide Complex Trait Analysis (GCTA) software.^21–23^ We report heritability on the observed scale due to genome-wide genotyped variants as well as heritability on the liability scale assuming AML and MDS disease prevalence of 0.0001.^24–26^

#### Transcriptome-wide association study (TWAS) of AML and MDS

To prioritize GWAS findings and identify expression quantitative trait loci (eQTL)-linked genes, we carried out a gene expression tests of association of *de novo* AML and MDS using PrediXcan^11^. This method leverages the well-described functional regulatory enrichment in genetic variants relatively close to the gene body (i.e. *cis*-regulatory variation) to inform models relating SNPs to gene expression levels in data with both gene expression and SNP genotypes available. Robust prediction models are then used to estimate the effect of cis-regulatory variation on gene expression levels. Using imputation, the cis-regulatory effects on gene expression from these models can be predicted in any study with genotype measurements, even if measured gene expression is not available. Thus, we imputed the cis-regulatory component of gene expression into our data for each individual using models trained on the whole blood transcriptome panel (n = 922) from the Depression Genes and Networks (DGN)^27^, yielding expression levels of 11,200 genes for each case and control. The resulting estimated gene expression levels were then used to perform gene-based tests of differential expression between AML and MDS cases and controls adjusted for age and sex. A fixed effects model with inverse variance weighting using the R package Metafor was used for meta-analysis of cohorts 1 and 2. A Bonferroni-corrected transcriptome wide significance threshold was set at P<4.5×10^-6^.

### Functional Annotation of Genetic Variation associated with AML and MDS

To better understand the potential function of the variants identified by GWAS and ASSET analyses we annotated significant SNPs using publicly available data. eQTLGen, a consortium analyses of the relationship of SNPs to gene expression in 30,912 whole blood samples, was used to determine if significant and suggestive SNPs (p<5 × 10^-6^) were whole blood *cis*-eQTL, defined as allele specific association with gene expression ^28^. Genotype-Tissue Expression project (GTEx) was used to test for significant eQTLs in >70 additional tissues ^29^. AML and MDS SNP associations were also placed in context of previous GWAS using Phenoscanner, a variant-phenotype comprehensive database of large GWAS, which includes results from the NHGRI-EBI GWAS catalogue, the UK Biobank, NIH Genome-Wide Repository of Associations between SNPs and Phenotypes and publicly available summary statistics from more than 150 published genome association studies. Results were filtered at P < 5 × 10^-8^ and the R statistical software package `phenoscanner` (https://github.com/phenoscanner/phenoscanner) was used to download all data for our significant variants^30^. Chromatin state data based on 25-state Imputation Based Chromatin State Model across 24 Blood, T-cell, HSC and B-cell lines was downloaded from the Roadmap Epigenomics project (https://egg2.wustl.edu/roadmap/data/byFileType/chromhmmSegmentations/ChmmModels/imputed12marks/jointModel/final/)^31^. Figures including chromatin state information and results from previous GWAS were constructed using the R Bioconductor package `gviz` ^32–34^. Lastly, we sought to identify promoter interaction regions (PIR), defined as significant interactions between gene promotors and distal genomic regions. Variants in PIRs can be connected to potential gene targets and thus can impact gene function ^34^. Briefly Hi-C libraries, enriched for promoter sequences, are generated with biotinylated RNA baits complementary to the ends of promoter-containing restriction fragments. Promoter fragments become bait for pieces of the genome that are targets with which they frequently interact, allowing regulatory elements and enhancers to be pulled down and sequenced. Statistical tests of bait-target pairs are done to define significant PIRs and their targets ^32^,^35^,^36^. To identify the genomic features with which our significant SNPs might be interacting via chromatin looping we used publicly available Promoter Capture Hi-C (PCHi-C) data on a lymphoblastoid cell line (LCL), GM12878, and two *ex vivo* CD34^+^ hematopoietic progenitor cell lines (primary hematopoietic G-CSF mobilized stem cells and hematopoietic stem cells) ^35^. We integrated our SNP data with the PCHi-C cell line data and visualized these interactions using circos plots ^37^.

## RESULTS

### DISCOVeRY-BMT cases and controls

Results of quality control have been described elsewhere.^14^ Following quality control, the DISCOVeRY-BMT cohorts include 1,769 AML and 540 MDS patients who received URD-BMT as treatment and 2,814 unrelated donors as controls (**Table 1**). The majority of AML cases are *de novo* (N=1618) with normal cytogenetics (N=543), 6% of patients had therapy-related AML (t-AML). The most frequently reported previous cancers in patients with t-AML were breast (N=51), non-Hodgkin Lymphoma (NHL), N=23, HL (N=14),

**Table 1.** DISCOVeRY-BMT Acute myeloid leukemia (AML) and myelodysplastic syndrome (MDS) Patient and Control Characteristics.

|  |  |  |
| --- | --- | --- |
| Patient and Donor Characteristics | Cases<br>Cohort 1 / Cohort 2<br>N= 1627 (%) / 682 (%) | Controls<br>Cohort 1 / Cohort 2<br>N= 2052 (%) / 762 (%) |
| **Age, years** |  |  |
| Median (range) | 50 (<1-74.5) / 52 (<1-78) | 33 (18-61) / 31 (18-60) |
| **Sex** |  |  |
| Females | 741 (46) / 312 (46) | 656 (32) / 209 (27) |
| **Disease** |  |  |
| **AML, all cases** | **1282 (79) / 487 (71)** | - |
| **de novo AML** | **1164 (72) / 454 (66)** | - |
| **de novo AML with normal cytogenetics** | **373 (23) / 170 (25)** | - |
| **de novo AML with abnormal cytogenetics** | **595 (37) / 241 (35)** | - |
| By Cytogenetic Subtype: |  |  |
| Core Binding Factor | 67 (11) / 32 (13) | - |
| MLL | 72 (12) / 48 (20) | - |
| Ph+ t(9;22) | 5 (1) / 1 (0) | - |
| APL t(15;17) | 18 (3) / 3 (1) | - |
| Any translocation | 97 (15) / 35 (15) | - |
| Trisomy 8 | 103 (17) / 22 (9) | - |
| Trisomy 13, 21 or 22 | 52 (9) / 24 (9) | - |
| del5/del7 | 123 (21) / 55 (23) | - |
| Any Trisomy | 195 (33) / 92 (38) | - |
| Any Monosomy | 153 (26) / 50 (21) | - |
| >3 cytogenetic abnormalities | 213 (36) / 88 (37) | - |
| **therapy-related AML** | **113 (7) / 33 (5)** | - |
| By Prior Diagnosis<sup>2</sup>: |  |  |
| Breast Cancer | 39 (35) / 12 (36) | - |
| Non-Hodgkin Lymphoma | 20 (18) / 3 (9) | - |
| Hodgkin Lymphoma | 11 (10) / 3 (9) | - |
| Sarcoma | 9 (3) / 8 (9) | - |
| Gynecologic Cancer | 6 (5) / 2 (6) | - |
| Acute Lymphoblastic Leukemia | 4 (4) / 2 (6) | - |
| Testicular Cancer | 4 (4) / 2 (6) | - |
| Other Disease | 20 (18) / 4 (12) | - |
| **MDS, all cases** | **345 (21) / 195 (29)** |  |
| **de novo MDS** | **294 (18) / 150 (22)** |  |
| By WHO subtype<sup>2</sup>: |  |  |
| MDS-unclassified<sup>3</sup> | 58 (17) / 35 (18) |  |
| RA, RA-RS | 91 (26) / 28(15) |  |
| RAEB-1, RAEB-2 | 153 (44) / 89 (46) |  |
| Chronic Myelomonocytic Leukemia | 42 (12) / 16 (8) |  |
| RCMD, RCMD-RS | 0 (0) / 25 (13) |  |
| **therapy-related MDS** | **51 (3) / 45 (7)** |  |
| By Prior Diagnosis<sup>2</sup>: |  |  |
| Non-Hodgkin Lymphoma | 15 (29) / 12 (27) | - |
| Breast Cancer | 8 (16) / 7 (16) | - |
| Hodgkin Lymphoma | 6 (12) / 2 (4) | - |
| Acute Lymphoblastic Leukemia | 4 (8) / 4 (9) | - |
| Acute Myeloid Leukemia | 4 (8) / 4 (9) | - |
| Chronic Lymphocytic Leukemia | 2 (4) / 3 (6) | - |
| Sarcoma | 1 (2) / 5 (11) | - |
| Other diseases | 10 (20) / 9 (20) | - |

<sup>1</sup>percentage of patient subgroup reflects the percentage of the total number of AML and MDS cases in each cohort; <sup>2</sup>percentage of patient subgroup reflects the percentage of the cases of corresponding disease subgroups in each cohort; <sup>3</sup>one individual had 5q-syndrome; RAEB=Refractory Anemia Excess Blasts; RCMD=Refractory Cytopenia with Multilineage Dysplasia; RCMD-RS=Refractory Cytopenia with Multilineage Dysplasia and Ringed Sideroblasts; RARS=Refractory Anemia with Ring Sideroblasts.

Sarcoma (N=12), Gynecologic (N=8), Acute Lymphoblastic Leukemia (N=6) and Testicular (N=6). Prior therapies for these patients were approximately equally divided between single agent chemotherapy and combined modality chemotherapy plus radiation. Almost half of MDS patients had Refractory Anemia with Excess Blasts (RAEB)-1 and RAEB-2. Of patients with t-MDS (~18% of MDS patients), 65% had antecedent hematologic cancers or disorders. The most frequently reported antecedent cancers in MDS patients were NHL (N=27), breast (N=15), Acute Lymphoblastic Leukemia (N=8), HL (N=8), AML (N=8), Sarcoma (N=6) and CLL (N=5) (**Table 1**). Overall, the distribution of antecedent cancers differed significantly between t-MDS and t-AML, with almost 2/3 of t-MDS and 1/3 of t-AML patients diagnosed with a prior hematologic cancer.

### SNP Associations with AML and MDS

GWAS of AML by subtype (abnormal cytogenetics, normal cytogenetics and t-AML) and MDS (*de novo* and t-MDS) are shown in **Supplemental Figure 1**. No population stratification was observed in PCA analysis and λ=1.0 in both cohorts.

To identify loci that show association with AML and MDS we used ASSET. For SNPs to be considered, we used previously defined criteria, which required ASSET SNP associations at P ≤ 5.0 × 10^-8^ with significant individual one-sided subset tests (P < 0.01), the variant association could not be driven by a single disease nor could it be both positively and negatively associated in different cohorts of the same disease.^5^ In the ASSET GWAS analyses we identified a novel typed SNP associated with AML and MDS on Chromosome 6 (**Figure 1**). The T allele at rs12203592, a variant in intron 4 of *Interferon Regulatory Factor 4 (IRF4)*, conferred increased risk of *de novo* abnormal cytogenetic AML, *de novo* normal cytogenetic AML, MDS and t-MDS (OR=1.38; 95% CI, 1.26-1.51, P_meta_=2.8 × 10^-12^). T-AML showed no association with rs12203592. The effect allele frequency was 19% in *de novo* AML, MDS and t-MDS cases versus 14% in controls. ASSET analyses also identified another variant in modest linkage disequilibrium (LD), r^2^=.7, with rs12203592 in the regulatory region of *IRF4;* the A allele at rs62389423, showed a putative association with de novo AML and MDS (OR=1.36; 95% CI, 1.21-1.52, P_meta_=1.2×10^-7^) (**Figure 2a**).

**Figure 1.**
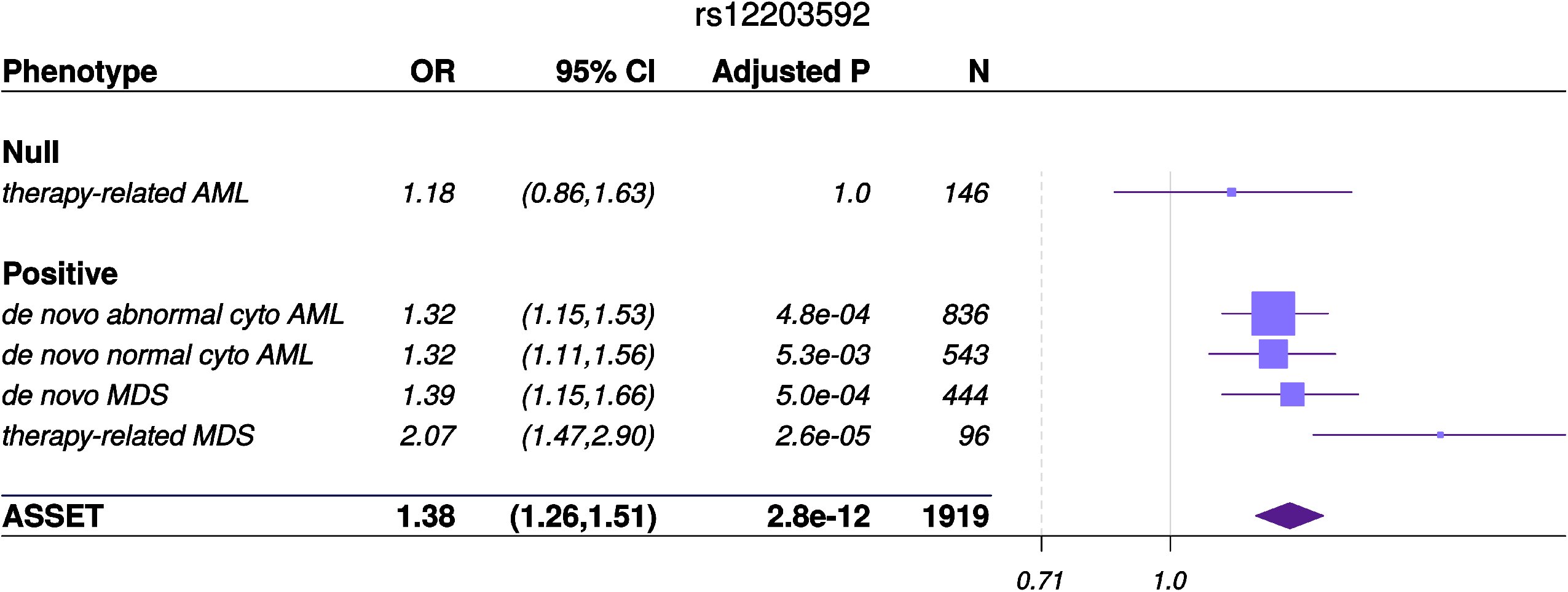
ASSET analysis and associations by AML and MDS subgroup. Forest plot of the odds ratios (OR) for the association between rs12203592 in *IRF4* and MDS and AML subtypes. The variant resides in the Chromosome 6 outside the major histocompatibility complex region. Studies were weighted by inverse of the variance of the log (OR). The solid grey vertical line is positioned at the null value (OR=1); values to the right represent risk increasing odds ratios. Horizontal lines show the 95% CI and the box is the OR point estimate for each casecontrol subset with its area proportional to the weight of the patient group. The diamond is the overall effect estimated by ASSET, with the 95% CI given by its width.

**Figure 2.**
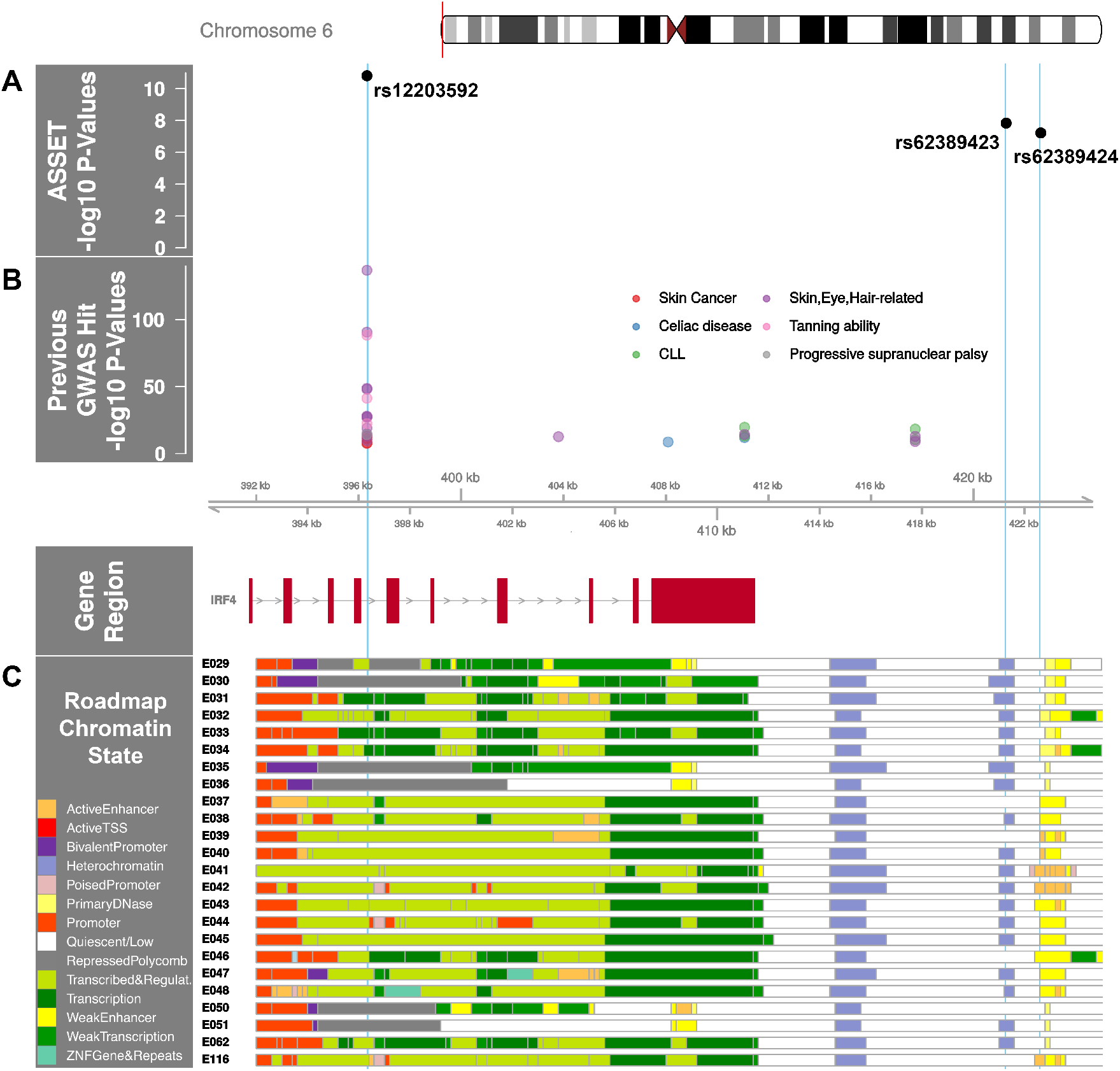
*IRF4* region with AML and MDS associated SNP p-values annotated with previous GWAS and Roadmap Epigenome Chromatin States. **A.** ASSET analysis AML and MDS SNP associations in the *IRF4* region. The x-axis is the chromosome position in kilobase pairs and y-axis shows the –log10 (p-values) for de novo AML and MDS susceptibility. The associated SNPs in the *IRF4* region, rs12203592 and rs62389423, are highlighted with sky blue lines drawn through the point to show the relationship of the variant to GWAS hits and Roadmap Epigenome data (2C). rs12203592 and rs62389423 show moderate linkage disequilibrium (r^2^=0.7); rs62389423 and rs62389424 are almost perfectly correlated (r^2^=.95). **B.** Previously reported GWAS SNPs in the *IRF4* region. Phenotypes are color coded and all variants are associated at P< 5 × 10^-8^. **C.** Genes in the region annotated with the chromatin-state segmentation track (ChromHMM) from Roadmap Epigenome data for all blood, T-cell, HSC and B-cells. The cell line numbers shown down the left side correspond to specific epigenome road map cell lines. E029:Primary monocytes from peripheral blood; E030:Primary neutrophils from peripheral blood; E031:Primary B cells from cord blood; E032:Primary B Cells from peripheral blood; E033:Primary T Cells from cord blood; E034:Primary T Cells from blood; E035:Primary hematopoietic stem cells; E036:Primary hematopoietic stem cells short term culture; E037:Primary T helper memory cells from peripheral blood 2; E038:Primary T help naïve cells from peripheral blood; E039:Primary T helper naïve cells from peripheral blood; E040:Primary T helper memory cells from peripheral blood 1; E041:Primary T helper cells PMA-Ionomycin stimulated; E042:Primary T helper 17 cells PMA-Ionomycin stimulated; E043:Primary T helper cells from peripheral blood; E044:Primary T regulatory cells from peripheral blood; E045:Primary T cells effector/memory enriched from peripheral blood; E046:Primary Natural Killer cells from peripheral blood; E047:Primary T CD8 naïve cells from peripheral blood; E048:Primary T CD8 memory cells from peripheral blood; E-50:Primary hematopoietic stem cells G-CSF mobilized Female; E-51:Primary hematopoietic stem cells G-CSF mobilized Male; E062:Primary Mononuclear Cells from Peripheral Blood; E0116 Lymphoblastic Cell Line. The colors indicate chromatin states imputed by ChromHMM and shown in the key titled “Roadmap Chromatin State”

We identified one significant association in the subtype GWAS which was disease specific. The C allele in rs78898975 in TATA-box binding protein associated factor 2 *(TAF2)*, associated with an increased risk of t-MDS (OR_meta_= 5.87, 95% CI = 3.20, 10.76, P_meta_=9.9 × 10^-9^) but not *de novo* MDS (OR= 1.8, 95% CI=.81, 1.45, P_meta_=.20) (**Supplemental Figure 1**). The effect allele frequency was 7% in t-MDS, 2% in *de novo* MDS and 1.5% in controls.

A previous genome-wide association study of AML done in European American cases and controls reported a susceptibility variant in *BICRA* (rs75797233) ^38^. The variant was not significantly associated with AML risk in our meta-analyses (OR=1.08, 95% CI=.78-1.37). However, their cohort did not include patients who received an allogeneic transplant as curative therapy and the distribution of AML subtypes differed between the studies. In addition, the lower frequency (MAF=.02) of this imputed this variant (info score >.8 in both cohorts) possibly reduced power to detect an effect.

### Functional Annotation of SNP associations with AML and MDS

Multiple GWAS of healthy individuals have shown associations between the T allele at rs12203592 and higher eosinophil counts, lighter skin color, lighter hair, less tanning ability, and increased freckling.^30,39^ GWAS have also identified associations between this allele and increased risk of childhood acute lymphoblastic leukemia in males, non-melanoma skin cancer, squamous cell carcinoma, cutaneous squamous cell carcinoma, basal cell carcinoma, actinic keratosis, and progressive supranuclear palsy (**Figure 2b**).^30^ Furthermore, analyses of multiple B-cell malignancies recently identified a rs9392017, adjacent to *IRF4*, as a pleiotropic susceptibility variant associated with both CLL and Hodgkin Lymphoma(HL) ^5^,^33^,^35^,^40^. This SNP is approximately 40Kb away from rs12203592, although not in LD (r^2^=.01).

The rs12203592 risk allele associated with increased expression of *IRF4*, P=1.48×10^-29^ in whole blood^28^. *IRF4* is a key transcription factor for lymphoid and myeloid hematopoiesis ^41–44^ and rs12203592 resides in a regulatory region across Blood, HSC, B-Cell and T-Cell lines (**Figure 2c**). The variant’s regulomedb score indicates how likely a variant is to be a regulatory element from 1a (most likely) to 7 (no data); the variant’s score of 2b, indicates the variant is likely to affect transcription factor binding^45^. While the HL and CLL pleiotropic variant rs9392017 was not a significant eQTL for *IRF4* in whole blood, PCHi-C cell line data from both GM12878 and the *ex vivo* CD34^+^ hematopoietic progenitor cell lines show chromatin looping between rs9392017 and the regulatory region containing rs12203592 (**Supplemental Figure 2**).

The t-MDS associated C allele in rs78898975 is correlated with significantly lower expression of *TAF2* (P=1.95 × 10^-13^) and *DEPTOR* (P= 4.7 × 10^-9^) gene expression in whole blood.^28,46^

### Heritability estimates of AML and MDS

The heritability of AML and MDS on the observed scale due to genotyped variants was 0.46 with standard error (SE)=0.07. Transforming this to the liability scale and assuming a disease prevalence of 0.0001 resulted in a heritability of 0.10 (SE=.02) which differed significantly from a heritability of zero (P=2.0 × 10^-16^). The proportion of variance in *de novo* AML with normal cytogenetics and *de novo* MDS on the liability scale had similar heritability at 9%, SE=.03, P=1.9 × 10^-3^ and 14%, SE=.04, P=1.4 × 10^-4^, respectively. Treatment-related AML and MDS were tested independently and estimated proportion of variance explained by all SNPs was 7% for t-AML and 4% for t-MDS, however SE were high and the heritability did not significantly differ from zero.

### Transcriptome-wide association study - PrediXcan

Using PrediXcan^11^ gene expression imputation models trained on the DGN data set, we identified one transcriptome wide significant gene associated with *de novo* AML and MDS. Increased expression of *IRF4* was associated with an increased risk for the development of *de novo* AML and MDS (OR=3.90; 95% CI, 2.36-6.44, P_meta_=1.0×10^-7^), consistent with our SNP-level findings (**Figure 3**).

**Figure 3.**
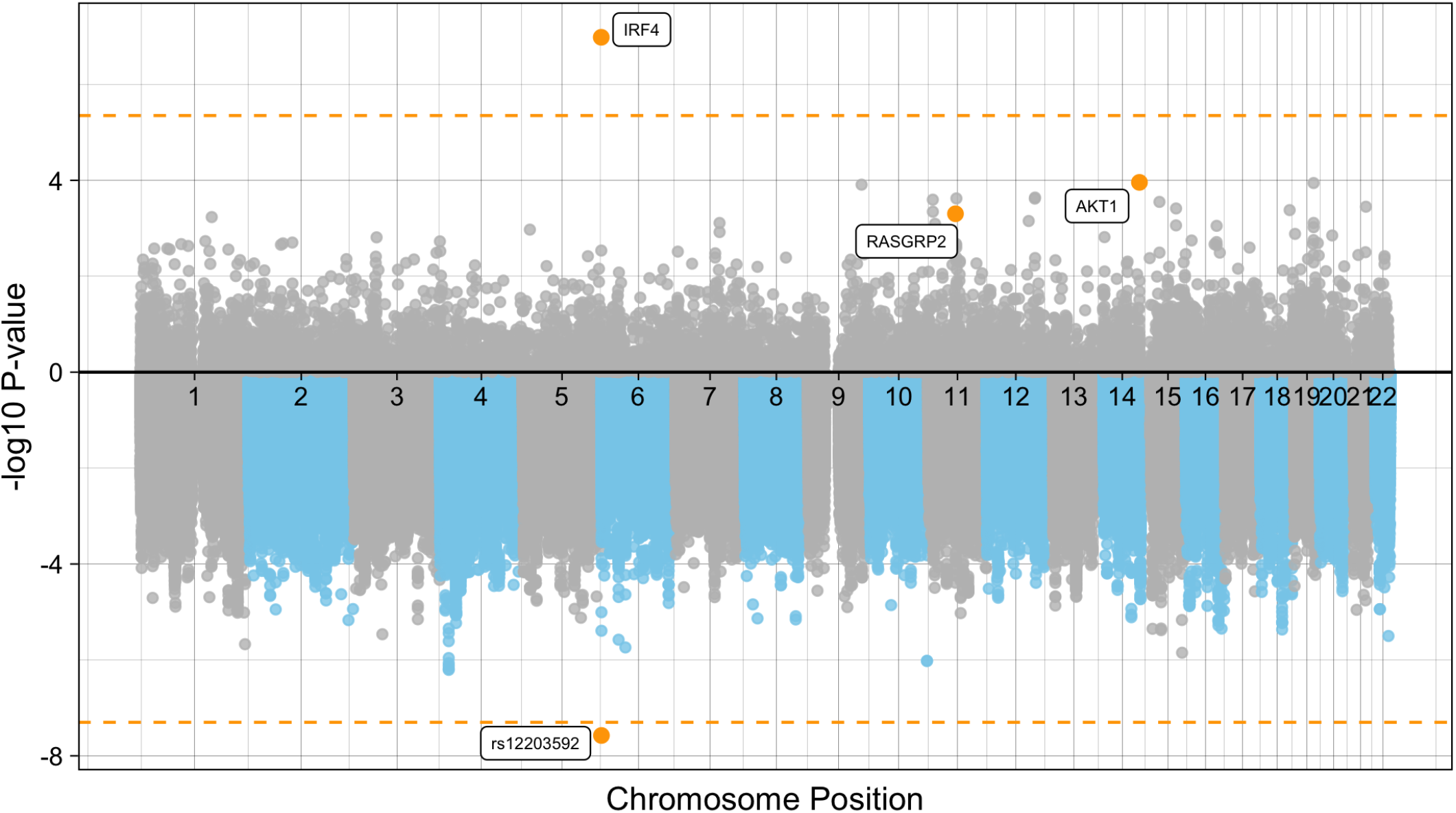
Manhattan plot of the *de novo* AML and MDS GWAS and TWAS. The plot represents the TWAS P-values (top) of each gene and *de novo* AML and MDS GWAS P-values (bottom) of each SNP included in the case-control association study. Significant and suggestive genes are highlighted in orange and labelled by their gene symbols. The orange horizontal line on the top represents the transcriptome-wide significance threshold of *P*=4.5×10^-6^. The orange horizontal line on the bottom represents the genome-wide threshold of *P*=5.0×10^-8^.

Whole blood transcriptome models also identified two additional genes with suggestive associations with *de novo* AML and MDS. Increased expression of AKT Serine/Threonine Kinase 1, *AKT1* at 14q32.33 was associated with risk for the development of *de novo* AML and MDS (OR=1.56; 95% CI, 1.25-1.95, P_meta_=1.0 ×10^-4^) (Figure 4). Likewise, increased expression of Ras guanyl nucleotide-releasing protein 2, *RASGRP2*, was associated with an increased risk for development of *de novo* AML and MDS (OR=4.05; 95% CI, 1.84-8.91, P_meta_=5×10^-4^) (**Figure 4**).

**Figure 4.**
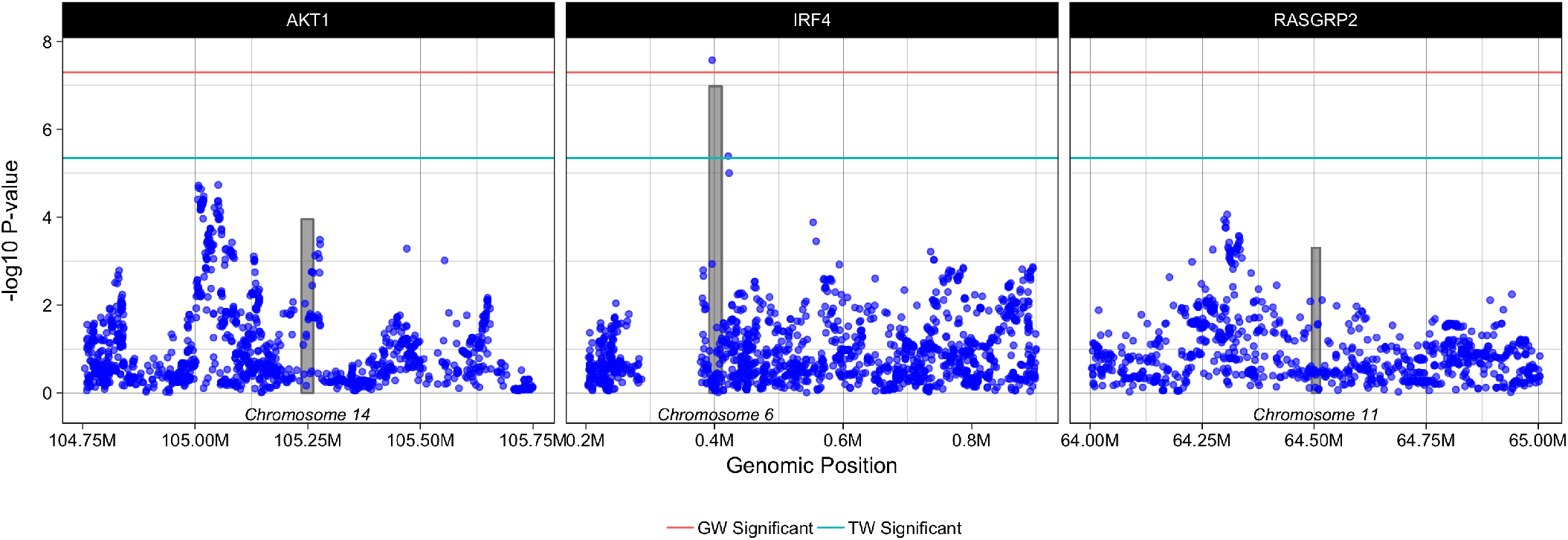
Regional plots of PrediXcan-TWAS and SNP associations with AML and MDS. Each box represents PrediXcan-TWAS significant genes *AKT1, IRF4* and *RASGRP2* +/-0.5 megabases. The grey shaded bars represent the gene, where height is gene expression association and width is gene region in base pairs and the purple dots represent SNP associations with AML and MDS -log10 (P-values) are shown on the y-axis. Green and red lines denote the transcriptome-wide and genome wide significant P-values, respectively.

## DISCUSSION

We performed the first large scale AML and MDS GWAS in a URD-BMT population providing evidence of novel pleotropic risk loci associated with increased susceptibility to AML and MDS. We identified an association between the T allele at rs12203592 in *IRF4* and an increased risk for the development of *de novo* AML, *de novo* MDS and t-MDS in patients who had undergone URD-BMT compared to healthy donor controls. While therapy-related myeloid neoplasms have been shown to be genetically and etiologically similar to other high-risk myeloid neoplasms^47^, in our transplant population t-AML did not associate with this variant, while t-MDS did show evidence of association with rs12203592. We also identified a genome-wide significant t-MDS variant which was an eQTL for both *TAF2* and *DEPTOR* genes. We also provide the first estimates of the heritability of AML and MDS, at between 9-14%, which are in line with other GWAS of cancer heritability on the liability scale, indicating that genetic variation contributes to AML and MDS susceptibility.^48^

The rs12203592 SNP has been shown to regulate *IRF4* transcription by physical interaction with the *IRF4* promoter through a chromatin loop^49^. This SNP resides in an important position within *NFkB* motifs in multiple blood and immune cell lines, supporting the hypothesis that this SNP may modulate *NFkB* repression of *IRF4* expression.^50^,^51^ Furthermore, this SNP resides in a hematopoietic transcription factor that has been previously identified to harbor a hematological cancer susceptibility locus, rs9392017, which we show interacts with the region containing our susceptibility variant. These data add to the mounting evidence that there could be pleiotropic genes across multiple hematologic cancers^5,52–55^.

Imputed gene expression logistic regression models showed a significant association between higher predicted levels of *IRF4* expression and the risk for development of *de novo* AML or MDS^11^. Although *IRF4* functions as a tumor suppressor gene in early B-cell development ^56^, in multiple myeloma *IRF4* is a well-established oncogene^44^, with oncogenic implications extending to adult leukemias^57^ and lymphomas^58^, as well as pediatric leukemia. *IRF4* overexpression is a hallmark of activated B-cell-like type of diffuse large B-cell lymphoma and associated with classical Hodgkin lymphoma (cHL), plasma cell myeloma and primary effusion lymphoma.^59^ In a case-control study of childhood leukemia increased *IRF4* expression was higher in immature B-common acute lymphoblastic leukemia and T-cell leukemia with the highest expression levels in pediatric AML patients compared to controls^60^. In addition to the CLL genetic susceptibility loci identified in *IRF4*, high expression levels of the gene have been shown to correlate with poor clinical prognosis ^61^.

TWAS studies can be a powerful tool to help prioritize potentially causal genes. It is, however, imperative to investigate the SNP and gene-expression associations in the context of the surrounding variants and genes to reduce the possibility of a false signal from co-regulation. Co-regulation can occur when there are multiple GWAS and TWAS hits due to linkage disequilibrium and thus it becomes difficult to determine which locus is driving the phenotypic association. In our study, the SNP rs12203592 is a significant eQTL for only *IRF4*, this implies that the SNP and imputed gene expression signal we identified is not being driven by co-regulation of neighboring SNPs and/or genes. When considering non-imputed gene expression sets, eQTLgen^28^ corroborates this finding; rs12230592 is significantly associated with only increased expression of *IRF4.* In addition, the relationship of rs12203592 to *IRF4* expression in blood seems tissue specific, as GTEx data across over 70 tissues shows association with only lung tissue at P=9.1×10^-9^. The specificity of rs12203592 to *IRF4* expression in blood and the lack of correlation between *IRF4* expression and other genes in DISCOVeRY-BMT give confidence that the observed ASSET association is the potential susceptibility locus in the region. The functional significance of variants in this gene in hematopoiesis and its previous recognition as a locus associated with the risk for development of other hematological malignancies, further strengthen the evidence of an association of IRF4 with development of AML and MDS.

In addition to *IRF4*, we identified an association between the risk for development of *de novo* AML or MDS and higher expression of *AKT1. AKT1* is an oncogene which plays a critical role in the *PI3K/AKT* pathway. AML patients frequently show increased *AKT1* activity, providing leukemic cells with growth and survival promoting signals^62^ and enhanced *AKT* activation has been implicated in the transformation from MDS to AML and overexpression of *AKT* has been shown to induce leukemia in mice.^63^

We also identified AML and MDS gene expression associations with *RASGRP2*, which is expressed in various blood cell lineages and platelets, acts on the *Ras*-related protein Rap and functions in platelet adhesion. GWAS have identified significant variants in this gene associated with immature dendritic cells (% CD32+) and immature fraction of reticulocytes, a blood cell measurement shown to be elevated in patients with MDS versus controls.^39^ *RASGRP2* expression has not been studied in relation to AML or MDS, however recently *RASGRP2/Rap1* signaling was shown to be functionally linked to the CD38-associated increased CLL cell migration. The migration of CLL cells into lymphoid tissues because of proliferation induced by B-cell receptor activation is thought to be an important component of CLL pathogenesis.^64^ This finding has implications for the design of novel treatments for CD38+ hematological diseases.^64^ These data imply the replication of these gene expression associations with the development of AML and MDS are warranted.

This is the largest genome-wide AML and MDS susceptibility study to date. Despite our relatively large sample size, the complexity of cytogenetic risk groups in these diseases limits our analysis, particularly with respect to therapy-related AML and MDS.

The DISCOVeRY-BMT study population is composed of mostly European American non-Hispanics and thus validation of these associations in a non-white cohort of patients is imperative. Lastly, the use of TWAS is a powerful way to start to prioritize causal genes for follow-up after GWAS, however there are limitations. TWAS tests for association with genetically predicted gene expression and not total gene expression, which includes environmental, technical and genetic components.^65^

Our results provide evidence for the impact of common variants on the risk for AML or MDS susceptibility and further characterization of the 6p25.3 locus might provide a more mechanistic basis for the pleiotropic role of *IRF4* in AML and MDS susceptibility. The co-identification of variants in *IRF4* associated with the risk for both myeloid and lymphoid malignancy supports the importance of broader studies that span the spectrum hematologic malignancies.

## ACKNOWLEDGEMENTS / CONTRIBUTIONS / FUNDING

This work was supported by grants from the National Institute of Health. LESC and TH were supported by 1R01HL102278 and 1R03CA188733 to perform this work. EK is supported by the Pelotonia Foundation Graduate Student Fellowship. Any opinions, findings, and conclusions expressed in this material are those of the author(s) and do not necessarily reflect those of the Pelotonia Fellowship Program or The Ohio State University.

ACG was supported by CA9204, Mayo Clinic R25 Training Grant when she performed a majority of this work. The CIBMTR is supported by Public Health Service Grant/Cooperative Agreement 5U24-CA076518 from the National Cancer Institute (NCI), the National Heart, Lung and Blood Institute (NHLBI) and the National Institute of Allergy and Infectious Diseases (NIAID); a Grant/Cooperative Agreement 5U10HL069294 from NHLBI and NCI; a contract HHSH250201200016C with Health Resources and Services Administration (HRSA/DHHS); two Grants N00014-15-1-0848 and N00014-16-1-2020 from the Office of Naval Research; and grants from Alexion; *Amgen, Inc.; Anonymous donation to the Medical College of Wisconsin; Astellas Pharma US; AstraZeneca; Be the Match Foundation; *Bluebird Bio, Inc.; *Bristol Myers Squibb Oncology; *Celgene Corporation; Cellular Dynamics International, Inc.; *Chimerix, Inc.; Fred Hutchinson Cancer Research Center; Gamida Cell Ltd.; Genentech, Inc.; Genzyme Corporation; *Gilead Sciences, Inc.; Health Research, Inc. Roswell Park Cancer Institute; HistoGenetics, Inc.; Incyte Corporation; Janssen Scientific Affairs, LLC; *Jazz Pharmaceuticals, Inc.; Jeff Gordon Children’s Foundation; The Leukemia & Lymphoma Society; Medac, GmbH; MedImmune; The Medical College of Wisconsin; *Merck & Co, Inc.; Mesoblast; MesoScale Diagnostics, Inc.; *Miltenyi Biotec, Inc.; National Marrow Donor Program; Neovii Biotech NA, Inc.; Novartis Pharmaceuticals Corporation; Onyx Pharmaceuticals; Optum Healthcare Solutions, Inc.; Otsuka America Pharmaceutical, Inc.; Otsuka Pharmaceutical Co, Ltd. – Japan; PCORI; Perkin Elmer, Inc.; Pfizer, Inc; *Sanofi US; *Seattle Genetics; *Spectrum Pharmaceuticals, Inc.; St. Baldrick’s Foundation; *Sunesis Pharmaceuticals, Inc.; Swedish Orphan Biovitrum, Inc.; Takeda Oncology; Telomere Diagnostics, Inc.; University of Minnesota; and *Wellpoint, Inc. The views expressed in this article do not reflect the official policy or position of the National Institute of Health, the Department of the Navy, the Department of Defense, Health Resources and Services Administration (HRSA) or any other agency of the U.S. Government.

*Corporate Members

## Authorship Contributions

J.W, A.C-G, L.S-C, and T.E.H designed the research, performed research and analysis, and wrote the manuscript.

C.A.H, D.V, X.S and L.P performed the genotyping.

X.Z., L.P, A.W and G.B performed quality control of genomic data.

All authors reviewed and approved the manuscript.

---

**Supplemental Figure 1.**
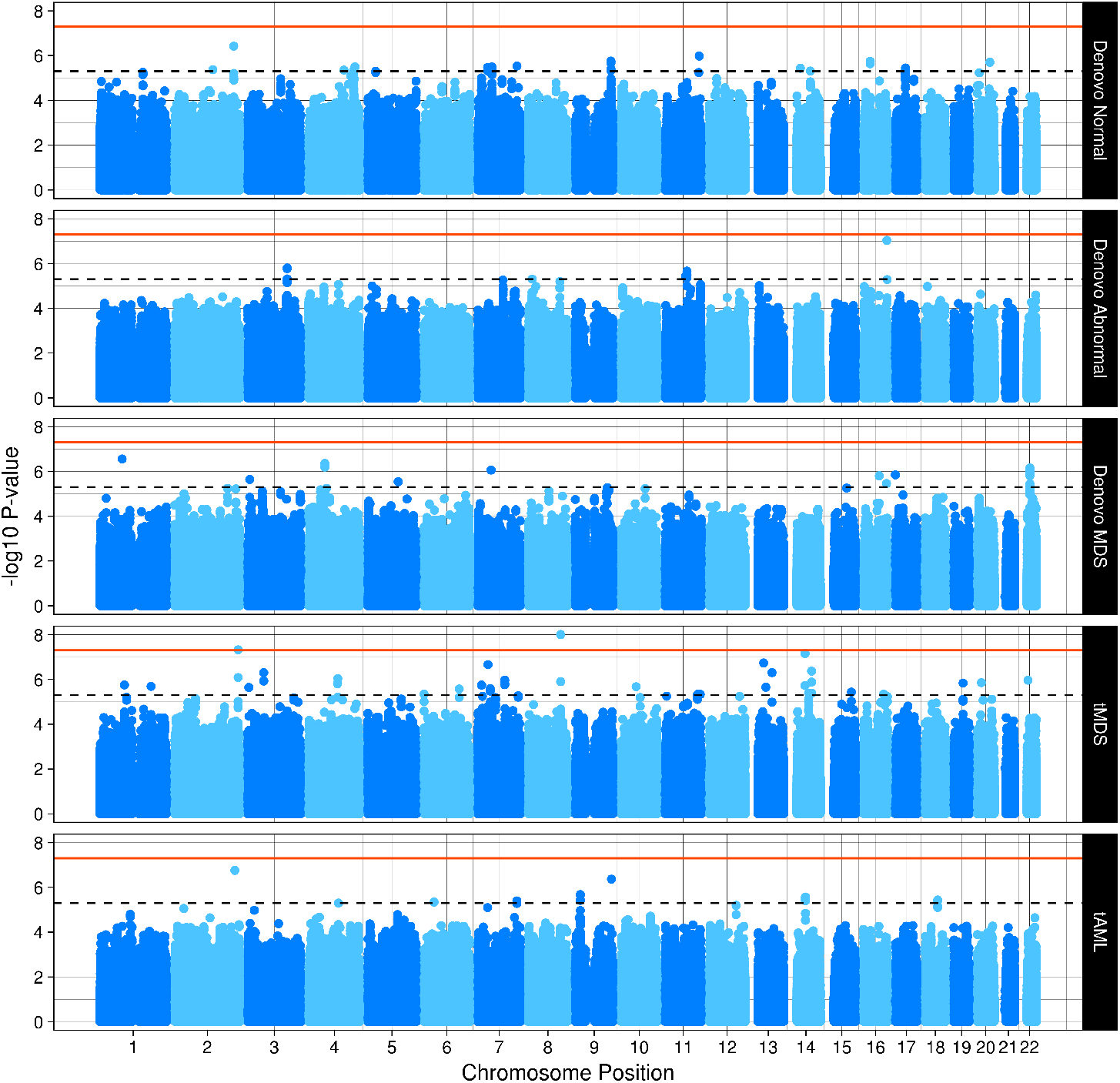
Genome wide associations by cytogenetic subtype in DISCOVeRY-BMT. Shown are the genome-wide *P* values by subtype from the meta-analysis of DISCOVeRY-BMT cohorts, including a total of 2158 AML and MDS cases and 2814 controls. The dashed horizontal line represents the suggestive threshold of *P*=5.0×10^-6^. The orange horizontal line represents the genome-wide significance threshold of *P*=5.0×10^-8^.

**Supplemental Figure 2.**
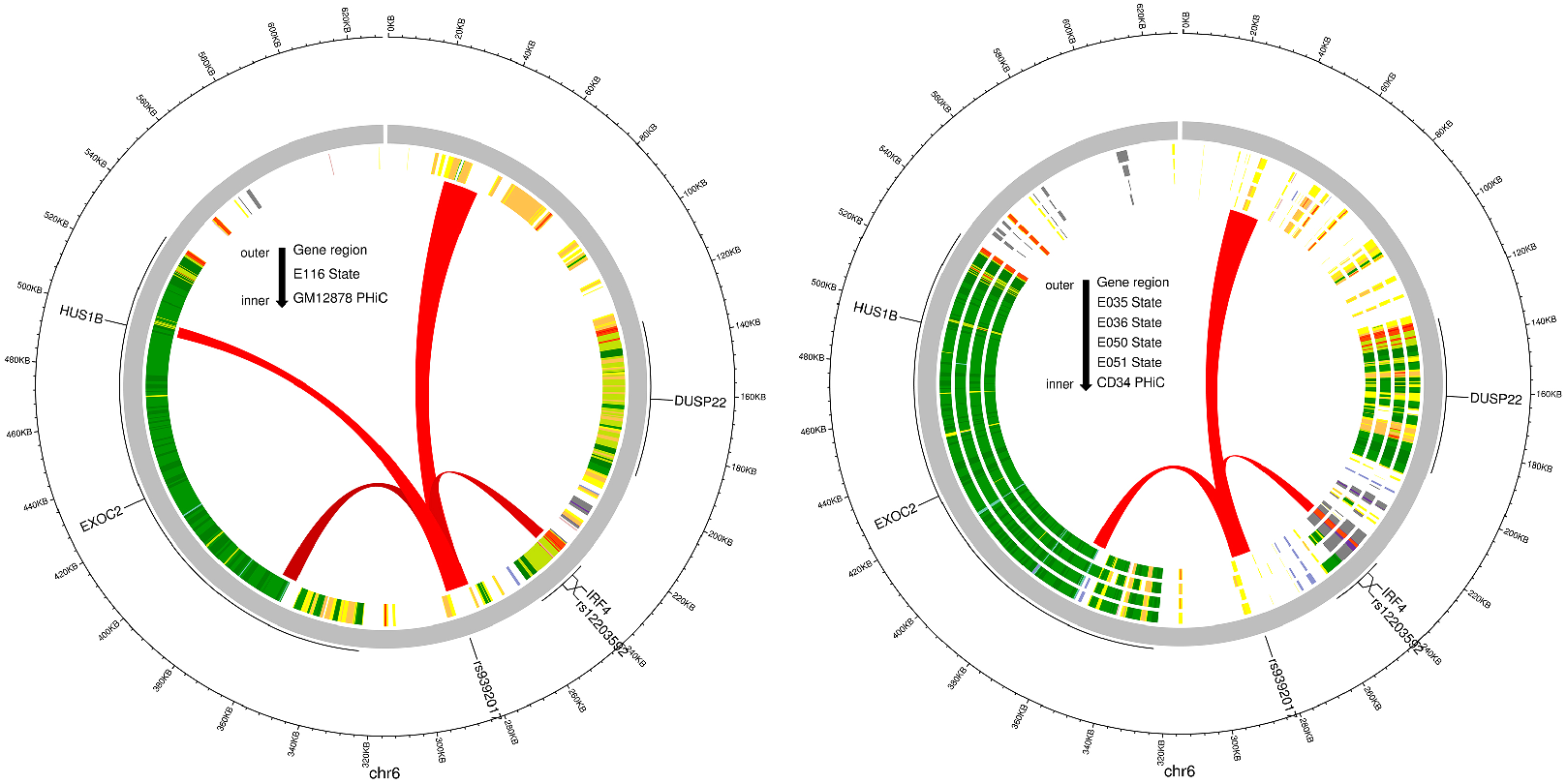
Significant chromatin interactions between the promoter region containing AML and MDS susceptibility variant, rs12203592 and the target region containing the previously identified CLL and HL susceptibility variant, rs9392017. The circular plots show significant chromatin interactions between bait-target pairs, defined as a CHICAGO score >=5, designated with red arcs, generated by promoter capture HI-C experiments in multiple cell lines. Moving from the outside of the circles inward we see base pair position on chromosome 6 in Kb, protein coding genes are shown in grey *(HUS1B, EXOC2, DUSP22* and *IRF4)*, the ENCODE roadmap epigenome chromatin states for **(LEFT)** E116: lymphoblastoid cell line and the following cell lines **(RIGHT)** E035:Primary hematopoietic stem cells; E036:Primary hematopoietic stem cells short term culture; E-50:Primary hematopoietic stem cells G-CSF mobilized Female; E-51:Primary hematopoietic stem cells G-CSF mobilized Male. This figure shows chromatin looping from the reference of the CLL and HL susceptibility region containing rs9392017 which illustrates this target region interacts with only few adjacent areas and only one transcriptional start site which contains rs12203592 providing support for the role of *IRF4* in CLL, HL, AML and MDS.

